# CRISPR-mediated transcriptional activation as a mutation-independent therapeutic strategy for *SYNGAP1*-related intellectual disability

**DOI:** 10.1101/2025.10.28.685100

**Authors:** Laura Sichlinger, Molly B. Reilly, Sakshi Arora, Shuo Zhang, Nicolas Marotta, Kiara L. Rodríguez-Acevedo, Marisol Hooks, Kyle S. Czarnecki, Julia J. Winter, Elisa A. Waxman, Lea V. Dungan, Ingie Hong, Yoichi Araki, Richard Johnson, Richard L. Huganir, Giulia Pavani, Deborah L. French, Beverly L. Davidson, Benjamin L. Prosser, Elizabeth A. Heller

## Abstract

Synaptic Ras GTPase-activating protein (SynGAP) regulates synaptic strength and neuronal signaling, with essential roles in cortical development and synaptic plasticity. Heterozygous loss-of-function variants in *SYNGAP1* cause *SYNGAP1*-related intellectual disability (SRID), a severe neurodevelopmental disorder characterized by epilepsy, developmental delay, and autism. *SYNGAP1* mutations often result in haploinsufficiency, providing a strong rationale for gene-targeted therapies. However, no treatment currently addresses the underlying genetic cause of SRID. Here, we developed a CRISPR-mediated transcriptional activation (CRISPRa) approach to upregulate the functional *Syngap1* allele in a SRID mouse model. CRISPRa activated *Syngap1*, normalized SynGAP protein expression and downstream signaling, and rescued working memory deficits. We validated the translational potential of this strategy in human induced pluripotent stem cell (hiPSC)-derived excitatory cortical neurons. CRISPRa rescued *SYNGAP1* in two distinct loss-of-function variant lines. Together, these findings demonstrate the feasibility of mutation-independent transcriptional activation as a therapeutic approach for SRID and its broader applicability to haploinsufficiency disorders.

## Main

Synaptic Ras GTPase-activating protein (SynGAP) is a highly abundant neuronal protein and essential for synaptic transmission^1,2^. It is predominantly localized to the postsynaptic density (PSD) of excitatory synapses in the cortex, hippocampus (HPC), striatum and olfactory bulb^3^. *SYNGAP1* is among the most frequently mutated genes in neurodevelopmental disorders (NDDs)^4^. Highly penetrant, heterozygous loss-of-function variants in *SYNGAP1* cause SynGAP haploinsufficiency, which leads to *SYNGAP1*-related intellectual disability (SRID)^5,6^. SRID is an NDD, and its symptoms follow a developmental trajectory^7^. Longitudinal clinical studies show global developmental delays and intellectual disability from early infancy, which usually precede the onset of epileptic seizures^7,8^. Approximately 80-85% of individuals with SRID have epilepsy with seizures typically starting between 2-3 years of age, though they can occur from 4 months to 7 years^7,9,10^. Additional symptoms include strabismus, motor deficits, autism spectrum disorder (ASD; in up to 50% of cases), sleep abnormalities, attentional problems, impulsivity, aggression, and sensory abnormalities such as hyperacusis, and elevated pain threshold^7,10–13^. Current treatments for SRID are limited to managing symptoms through physical and behavioral therapies or anti-seizure medications^7^. No therapies yet exist to treat underlying genetic mutations or protein haploinsufficiency.

SynGAP is a multifunctional protein. It is a negative regulator of Ras signaling by promoting GTP hydrolysis to GDP through its GAP domain^1,2,14^ and is essential for activity-dependent synaptic plasticity^15,16^. At basal levels, SynGAP binds to membrane-associated guanylate kinase (MAGUK) proteins such as PSD-95^17^ and inhibits Ras-Raf-MEK-ERK signaling involved in long-term potentiation (LTP)^18–20^. Upon synaptic stimulation, SynGAP is phosphorylated by Ca^2+^/calmodulin-dependent protein kinase II (CaMKII) and translocates away from the PSD^15,21^. This was shown to relieve its inhibitory effects and trigger downstream cellular processes, such as GTPase activation, AMPA receptor insertion, and dendritic spine enlargement^15,18^. A recent study using catalytically inactive SynGAP constructs *in vitro* and a mouse model with inactivating GAP mutations showed that the GAP activity of SynGAP is not required for LTP or related cognitive functions^22^. Instead, it revealed a distinct structural role for SynGAP within the PSD, where it undergoes liquid-liquid phase separation with PSD-95 and competes with AMPAR/transmembrane AMPA receptor regulatory protein (TARP) complexes for PDZ-binding sites^22,23^. Beyond the synapse, SynGAP is critical for early cortical development. *SYNGAP1* haploinsufficiency in human induced pluripotent stem cell (hiPSC)-derived cortical organoids disrupted apico-basal polarity in neural progenitors, disorganized cortical plate formation, and accelerated neuronal maturation^24^. *SYNGAP1*-deficient postmitotic neurons exhibited increased dendritic growth^25^. These studies highlight functions for *SYNGAP1* in both neurodevelopmental and mature synaptic function.

Heterozygous *Syngap1* knockout mice recapitulate key SRID endophenotypes, including deficits in learning, memory, social behavior, as well as hyperactivity, and seizures^26–28^. Furthermore, *Syngap1* haploinsufficiency in mice impairs tactile processing through reduced connectivity and excitability of upper-lamina somatosensory neurons^29^ and disrupts cortical-thalamic dynamics and long-range connectivity^30^. This leads to deficits in tactile sensitivity and sensorimotor integration, highlighting circuit-specific pathologies that may contribute to the complex sensory and cognitive phenotypes of SRID. To investigate mechanisms specific to SRID-causing genetic mutations, knock-in mouse models carrying patient-specific *SYNGAP1* variants were generated^31^. These mouse models also recapitulate SRID endophenotypes and represent a genetic environment that most resembles patient cells. Hence, they provide a suitable preclinical platform for the evaluation of mechanistic consequences of pathogenic variants and for developing targeted treatments for SRID.

Recent advances in gene engineering technologies based on clustered regularly interspaced short palindromic repeats (CRISPR)-CRISPR-associated protein 9 (Cas9) demonstrated the therapeutic potential of genome editing for human genetic disorders. FDA-approved CRISPR-based therapeutics and ongoing clinical trials have already improved outcomes for patients with sickle cell disease, β-thalassemia, and hereditary angioedema^32–34^. Moreover, a recent *n = 1* clinical trial showed the feasibility of personalized, corrective base-editing for ultrarare variants, highlighting the precision and adaptability of CRISPR-based medicine^35^. However, traditional approaches rely on DNA double-strand breaks and sequence-specific corrections, which limits their potential use for genetically heterogeneous disorders. In contrast, CRISPR-mediated transcriptional modulation uses a catalytically dead Cas9 protein (dCas9) fused to transcriptional regulators to manipulate endogenous gene expression without altering the DNA sequence^36^. This strategy is reversible, minimizes off-target effects, and is not mutation-specific. Hence, it is promising for treating SRID.

Here, we developed a CRISPR-mediated transcriptional activation (CRISPRa) strategy that activates *Syngap1* expression in a preclinical SRID mouse model^31^, restoring ERK signaling and rescuing an SRID-associated behavior. We further demonstrated the translatability of this approach by activating human *SYNGAP1* in excitatory cortical neurons from patient-derived hiPSCs with distinct pathogenic *SYNGAP1* mutations, supporting its potential applicability across different SRID-causing variants.

## Results

### c.3583-9 G>A mutation leads to *Syngap1* haploinsufficiency

To evaluate the therapeutic potential of *Syngap1* CRISPRa, we selected a previously validated knock-in mouse model, *Syngap1^+/c.3583–9G>A^*^31^. *Syngap1^+/c.3583-9G>A^* mice carry a heterozygous SRID patient-associated intronic mutation (c.3583-9G>A) that generates a cryptic splice site upstream of exon 17 of *Syngap1*^37^. This leads to aberrant splicing and the inclusion of 7 base pairs (bp) in front of exon 17 within mRNA transcripts, which in turn leads to a premature stop codon (**Fig. 1A**). First, we confirmed the effect of this mutation on *Syngap1* expression by extracting total mRNA from two 2 mm punches of the nucleus accumbens (NAc), prefrontal cortex (PFC), and dorsal hippocampus (dHPC) of adult male and female mutant and control mice. RT-qPCR analysis revealed that *Syngap1* expression varied significantly by genotype and brain region, but not by sex. Specifically, *Syngap1* mRNA levels were significantly reduced in *Syngap1^+/c.3583-9G>A^* mice across all regions, with an average reduction of 42.9 ± 16% in the NAc, 30 ± 26% in the PFC, and 45.5 ± 29% in the HPC compared to controls (**Fig. 1B**). We next confirmed the presence of the mutant transcript containing the 7bp insertion upstream of exon 17 in HPC samples. To specifically detect the mutant transcript, we designed a RT-qPCR assay with a reverse primer spanning the exon 16-17 junction including the additional 7bp at the start of exon 17 (**Fig. 1C**). We detected the mutant transcript only in *Syngap1^+/c.3583-9G>A^* mice, not in controls (**Fig. 1D**). Moreover, expression of the mutant transcript negatively correlated with overall *Syngap1* mRNA levels, suggesting it contributes to the observed *Syngap1* reduction (**Fig. 1E**). Interestingly, we noticed that *Syngap1* mRNA expression in control mice showed greater variability than in *Syngap1^+/c.3583-9G>A^* mice (**Fig. 1B**), which was confirmed by Bartlett’s test indicating a difference in variance (χ^2^(1) = 3.66, p = 0.056) across genotype groups (**Fig. 1F**). At the protein level, western blot analysis confirmed reduced SynGAP protein expression in NAc, PFC, and HPC of *Syngap1^+/c.3583-9G>A^* mice with significant effects of genotype and region on SynGAP protein expression (**Fig. 1G & H**). While region-specific differences were not statistically significant in post-hoc comparisons, *Syngap1^+/c.3583-9G>A^*mice consistently showed reduced SynGAP protein: 25% in the NAc, 40% in the PFC, and 40% in the HPC (**Fig. 1H**). Together, these findings are in line with previous literature^31^ and support that *Syngap1^+/c.3583-9G>A^* mRNA is likely targeted by nonsense-mediated decay (NMD). Hence, this mouse model provides a suitable platform to test CRISPRa strategies for rescuing *Syngap1* haploinsufficiency.

**Figure 1.**
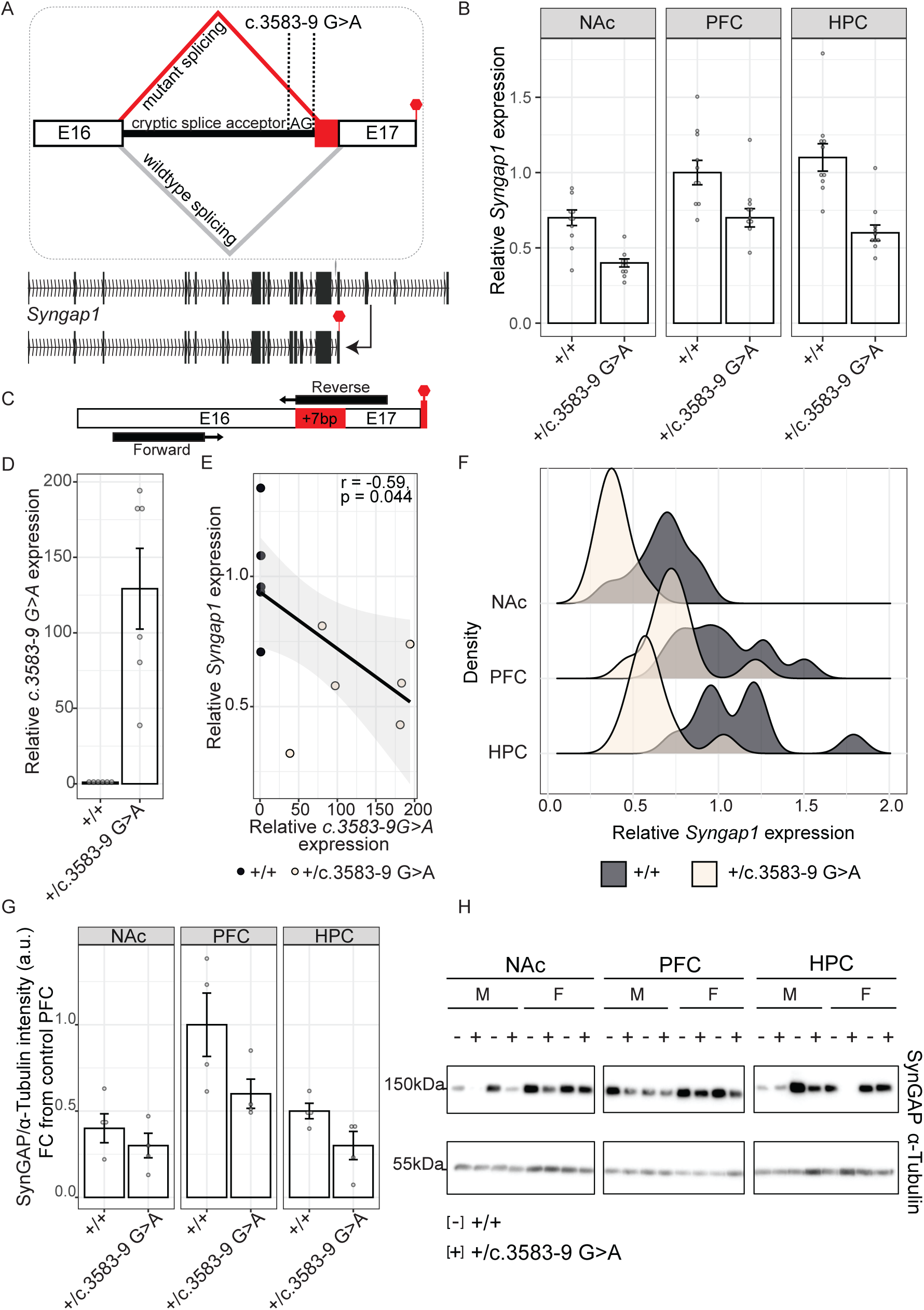
c.3583-9 G>A mutation leads to *Syngap1* haploinsufficiency. (A) Illustration of the *Syngap1* intronic mutation (c.3583-9G>A) upstream of exon 17, introducing a cryptic splice acceptor, which leads to aberrant splicing, inclusion of 7 bp, and a premature stop codon. (B) *Syngap1* mRNA expression (RT-qPCR) in two 2 mm punches from the nucleus accumbens (NAc), prefrontal cortex (PFC), and hippocampus (HPC) of adult *Syngap1^+/c.3583-9G>A^*and control mice (n = 10 per group; control: 7F/3M, *Syngap1*^+/c.3583-9G>A^: 5F/5M). Three-way ANOVA revealed significant effects of genotype (*F*(1,53) = 45.30, *p* = 1.23 x 10^-8^) and region (*F*(2,53) = 19.77, *p* = 3.86 x 10^-7^), but not sex (*F*(1,53) = 1.00, *p* = 0.322). TukeyHSD showed *Syngap1* expression was lower in NAc vs. HPC (p = 6.9 x 10^-6^) and vs. PFC (*p* = 2.4 x 10^-6^); no difference between HPC and PFC. *Syngap1^+/c.3583-9G>A^*mice showed reduced expression within regions: 42.9 ± 16% in NAc (TukeyHSD: *p* = 0.034), 30 ± 26% in PFC (TukeyHSD: *p* = 0.036), and 45.5 ± 29% in HPC (TukeyHSD: *p* = 1.8 x 10^-5^). (C) Schematic of c.3583-9G>A mutant specific primer pair. (D) RT-qPCR detection of *Syngap1* c.3583-9G>A mutant transcript in HPC of adult male and female mice (n = 6 per group; control: 4F/2M, *Syngap1*^+/c.3583-9G>A^: 1F/5M). Results only showed amplification of mutant transcript in *Syngap1^+/c.3583-9G>A^* mice not in controls (Wilcoxon rank sum test: W = 36, p = 0.002). (E) Scatter plot showing a significant negative correlation between expression of the mutant *Syngap1^+/c.3583-9G>A^* transcript and total *Syngap1* mRNA levels in HPC of adult mice (Pearson’s *r* = −0.59, *t*(10) = −2.30, *p* = 0.044). (F) Density plot showing *Syngap1* mRNA expression variability across brain regions (NAc, HPC and PFC). Bartlett’s test indicates a trending difference in variance between genotypes (*χ^2^*(1) = 3.66, *p* = 0.056), with *Syngap1^+/c.3583-9G>A^* mice exhibiting reduced expression variability across brain regions. (G) SynGAP protein expression in NAc, PFC, and HPC of adult *Syngap1^+/c.3583-9G>A^*and control mice (n = 4 per group; control: 2F/2M, *Syngap1*^+/c.3583-9G>A^: 2F/2M), normalized to loading control α-Tubulin. Two-way ANOVA of staining intensity ratios revealed significant main effects of genotype (*F*(1,18) = 9.50, *p* = 0.006) and brain region (*F*(2,18) = 11.83, *p* = 0.0005). While TukeyHSD post-hoc tests did not identify region-specific differences, *Syngap1^+/c.3583-9G>A^*mice showed reduced SynGAP protein across all regions: 25 ± 5.8% in NAc, 40 ± 19.8% in PFC, and 40 ± 14.6% in HPC. (H) Representative western blot of SynGAP protein (∼150kDa) levels in NAc, PFC, and HPC in *Syngap1^+/c.3583-9G>A^*(+) and control (−) mice. Loading control α-Tubulin (∼55kDa) probed on the same membrane shown on the bottom. Bars represent mean ± SEM error bars.

### CRISPR-mediated transcriptional activation strategy activates *Syngap1* in +/c.3583-9 G>A primary cortical neurons

Having validated *Syngap1* haploinsufficiency, we next developed a CRISPRa system to test whether *Syngap1* gene induction can restore *Syngap1* expression levels. We first trialed a CRISPRa system that we previously demonstrated to reliably induce gene expression *in vivo*^36^. The initial system consisted of a catalytically inactive Cas9 protein fused to the VP64 transcriptional activation domain (dCas9-VP64), driven by a neuron-specific promoter, along with a single-guide RNA (gRNA) targeting the *Syngap1* promoter (**Extended Data Fig. 1A**). For control experiments, we combined dCas9-VP64 with a non-targeting (nt) gRNA that has no sequence homology to the mouse genome (**Supplementary Table 1**). Using this system, we screened a library of gRNAs targeted at the *Syngap1* promoter up-stream (−891, –831, 574, –428, –346, –182, –18 bp) or down-stream (+351 bp) of the transcription start site (TSS) in Neuro2a (N2a) cells by JETPEI transfection (**Extended Data Fig. 1B & Supplementary Table 1**). In N2a cells, none of the tested gRNAs activated *Syngap1* mRNA expression (**Extended Data Fig. 1C-E**). We therefore revised our *Syngap1* CRISPRa approach.

We next adapted a lentivirus-compatible dual-component system previously optimized for neuronal target genes^38^. This system consisted of (1) a strong tripartite transcriptional activator (VP64, p65, and Rta) fused to a dCas9 protein (dCas9-VPR), driven by the neuron-specific promotor hSyn and (2) a gRNA targeting the *Syngap1* promotor region co-expressed with mCherry (**Fig. 2A**). To identify the most effective site for *Syngap1* activation, we designed a library of gRNAs targeted to a region of permissive histone post-translational modifications (hPTM) H3K4me3 (histone H3 lysine 4 tri-methylation) enrichment near the *Syngap1* TSS (**Fig. 2B**)^39^. The gRNAs targeted the dCas9-VPR system to sites both up-stream (−572, –288 bp) and down-stream (+2, +44 bp) of the TSS (**Fig. 2C**). After packaging each construct into lentiviral vectors, we initially screened the gRNA library in N2a cells, using a non-targeting gRNA with no homology to the mouse genome combined with dCas9-VPR as a control (nt control). Three days post-treatment, Syngap1 g44 activated *Syngap1* mRNA (**Fig. 2D-E**). However, *Syngap1* mRNA expression showed high variability across treatment conditions compared to controls, possibly due to the low baseline expression of *Syngap1* in N2a cells. Therefore, we proceeded to screen the entire CRISPRa library including the nt control and a dCas9-VPR construct targeted at an unrelated gene locus (Cartpt g-205) as a second control in a model system with more stable *Syngap1* expression. We dissected primary cortical neurons from *Syngap1^+/c.3583-9G>A^* embryos and control littermates, started lentiviral vector CRISPRa treatment at day in vitro (DIV) 4 and measured mRNA expression seven days later at DIV11 (**Fig. 2F**). We confirmed *Syngap1* mRNA activation by Syngap1 g44-dCas9-VPR in *Syngap1^+/c.3583-9G>A^* cortical neurons. Interestingly, Syngap1 g-288 only activated *Syngap1* mRNA in *Syngap1^+/c.3583-9G>A^* mice, not in controls (**Fig. 2G**). We concluded that Syngap1 g44-dCas9-VPR induced *Syngap1* mRNA expression in *Syngap1^+/c.3583-9G>A^* primary cortical neurons, supporting its further investigation *in vivo*.

**Figure 2.**
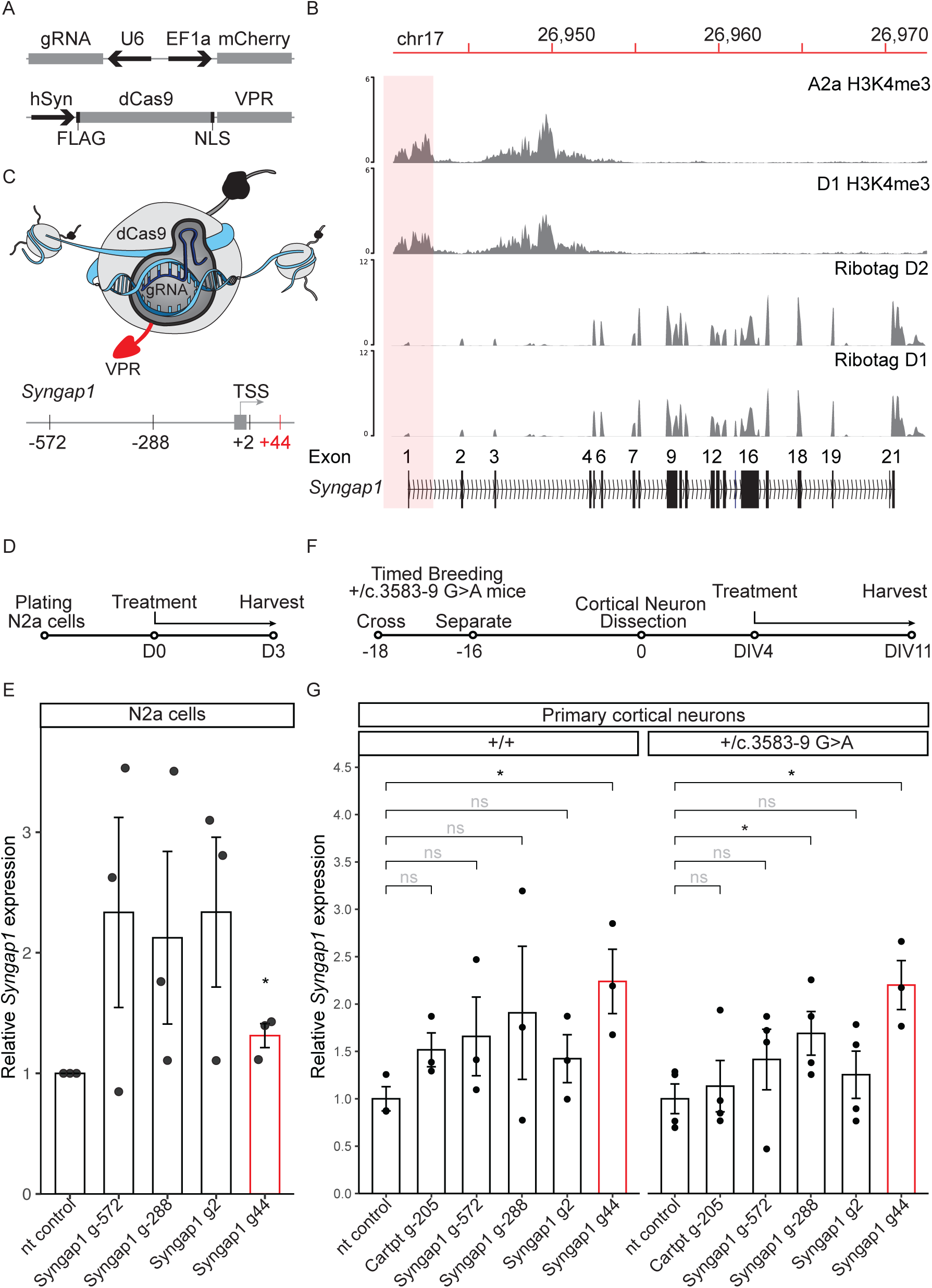
CRISPR-mediated transcriptional activation strategy up-regulates *Syngap1* in +/c.3583-9 G>A primary cortical neurons. (A) Illustration of the dual lentiviral vector CRISPR activation (CRISPRa) approach expressing (1) guide RNA (gRNA)-mCherry construct driven by U6 and EF1a promotors and (2) the dCas9-VPR construct driven by human Syn (hSyn) promoter. Adapted from^38^. (B) Genome browser view of gene activation-associated histone mark H3K4me3 enrichment near *Syngap1* transcription start site (TSS) in striatal neurons (A2a and D1). Tracks are visualized by pyGenomeTracks^63^. Data from^39^. H3K4me3 profiles were integrated with D1– and D2-specific translatosome data (RiboTag RNA profiling)^64^ to identify the optimal region to target for transcriptional activation of *Syngap1*. The red box highlights the locus targeted for *Syngap1* CRISPRa gRNA design. (C) Illustration of CRISPRa tool and gRNA library up-(−572 and –288 base pairs (bp)) and down-stream (2 and 44 bp) of the TSS of mouse *Syngap1*. (D) Timeline of lentiviral vector treatment of N2a cells. (E) *Syngap1* mRNA levels (RT-qPCR) in N2a cells (n = 3) treated with lentiviral vectors expressing CRISPRa non-targeting (nt) control and targeted at *Syngap1* locus (Syngap1 g-572, g-288, g2 and g44). *Syngap1* mRNA expression is shown relative to nt control within same plate. Two-tailed t-test showed significant up-regulation of Syngap1 g44-dCas9-VPR compared to nt control (t(4) = 3.13, *p* = 0.0350). (F) Timeline of lentiviral vector treatment of primary cortical neurons from *Syngap1^+/c.3583–9G>A^* embryos and litter controls. DIV = Day *in vitro*. (G) *Syngap1* mRNA expression (RT-qPCR) in primary cortical neurons from *Syngap1^+/c.3583-9G>A^* (n = 4) and control (n = 3) embryos treated with lentiviral vectors expressing nt control dCas9-VPR and targeted at *Cartpt* (Cartpt-205) and *Syngap1* locus (Syngap1 g-572, g-288, g2 and g44). Two-tailed t-test showed significant up-regulation of Syngap1 g44-dCas9-VPR (red) compared to nt control in both *Syngap1^+/c.358-9G>A^* (t(5) = 4.21, *p* = 0.008) and control (t(4) = 3.41, *p* = 0.027) cortical neurons. Syngap1 g-288-dCas9-VPR only showed *Syngap1* activation in *Syngap1^+/c.3583-9G>A^* neurons (t(6) = 2.48, *p* = 0.048). Bars represent mean ± SEM error bars, * p < 0.05.

### Syngap1 CRISPRa activates *Syngap1* in +/c.3583-9 G>A mouse hippocampus

To investigate whether Syngap1 CRISPRa is a viable therapeutic strategy for *Syngap1* haploinsufficiency, we aimed to rescue *Syngap1* haploinsufficiency *in vivo* in the *Syngap1^+/c.3583-9G>A^* mouse model using CRISPR-mediated *Syngap1* gene induction. We selected the HPC as the target region based on three considerations: (1) *Syngap1* is endogenously highly expressed in the HPC^40,41^, making it a biologically relevant and tractable site for intervention. (2) It showed the most pronounced reduction in *Syngap1* expression in our analyses (**Fig. 1B & G**) and, (3) HPC-dependent memory deficits have been reported in *Syngap1* heterozygous knock-out mouse models^28^ and *Syngap1^+/c.3583-9G>A^* mice^31^. Because Syngap1 g44-dCas9-VPR showed activation of *Syngap1* in +/c.3583-9 G>A primary cortical neurons, we chose to deliver the dual-lentiviral Syngap1 g44-dCas9-VPR system to the HPC of *Syngap1^+/c.3583-9G>A^*mice by stereotaxic surgery. In our stereotaxic transductions, expression is predominantly localized to the dHPC, with only minimal expression observed in ventral regions. In all experiments, Syngap1 g44-dCas9-VPR and the nt control were injected into opposite hemispheres, enabling rigorous within-animal comparisons (**Fig. 3A**). By qPCR, we confirmed activation of *Syngap1* mRNA by Syngap1 g44-dCas9-VPR relative to nt control dCas9-VPR in HPC of *Syngap1^+/c.3583-9G>A^*mice 14 days after surgery. Interestingly, Syngap1 g44-dCas9-VPR did not upregulate *Syngap1* mRNA in control mice (**Fig. 3B**), consistent with previous reports of CRISPRa targeting other dosage-sensitive genes^42,43^. To investigate if *Syngap1* activation results in increased SynGAP protein, we extracted protein from HPC tissue punches and analyzed protein expression by western blot. Syngap1 g44-dCas9-VPR upregulated SynGAP protein in *Syngap1^+/c.3583-9G>A^*HPC, but not in controls (**Fig. 3D**).

**Figure 3.**
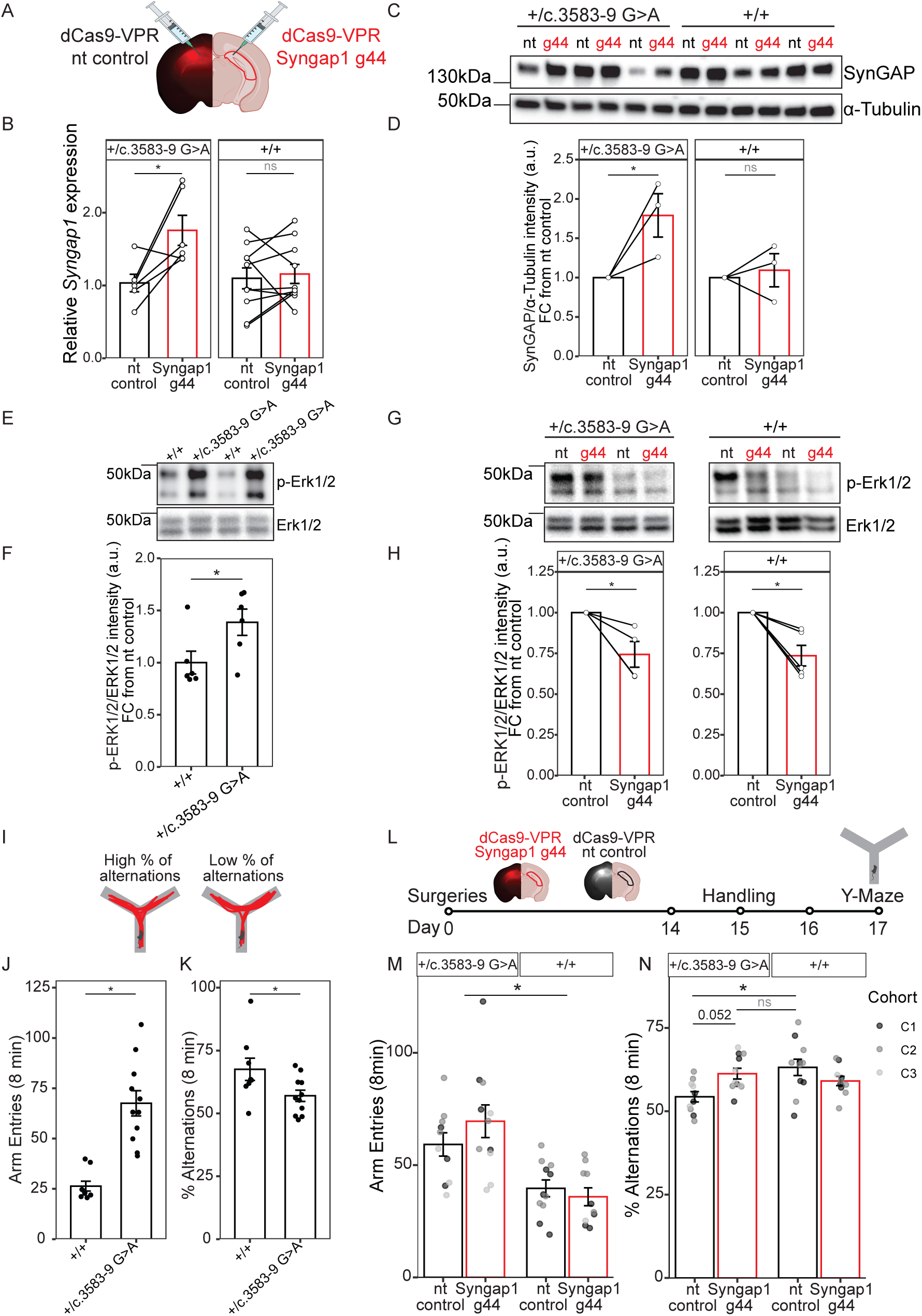
Syngap1 CRISPRa activates *Syngap1* in *+/c.3583-9 G>A* mouse hippocampus and rescues SRID-related behavior. (A) Illustration of contralateral stereotaxic delivery of dual lentiviral CRISPRa vectors, with *Syngap1* g44-dCas9-VPR injected into one hippocampal hemisphere and non-targeting (nt) control dCas9-VPR into the opposite hemisphere of the same mouse. (B) *Syngap1* mRNA expression (RT-qPCR) in mouse hippocampus (HPC) after injection of lentiviral vectors expressing either CRISPRa nt control dCas9-VPR or Syngap1 g44-dCas9-VPR (red). *Syngap1* mRNA expression is shown relative to nt control. Paired t-test showed significant up-regulation of *Syngap1* after Syngap1 g44-dCas9-VPR compared to nt control dCas9-VPR in *Syngap1^+/c.3583-9G>A^* mice (t(5) = –3.03, *p* = 0. 0288, n = 6, left) but not controls (t(9) = –0.46, *p* = 0.6562, n = 10, right). (C) Representative Western blot of SynGAP protein (∼150kDa) levels in HPC in *Syngap1^+/c.3583-9G>A^* and control mice, which received contralateral injections of either nt control dCas9-VPR (nt) or Syngap1 g44-dCas9-VPR (g44 in red). Loading control α-Tubulin (∼55kDa) probed on the same membrane (bottom). (D) SynGAP protein expression in mouse HPC after injection of lentiviral vectors expressing either CRISPRa nt control dCas9-VPR or Syngap1 g44-dCas9-VPR (red). SynGAP expression is shown relative to nt control within same animal. Paired t-test showed significant up-regulation of SynGAP after Syngap1 g44-dCas9-VPR compared to nt control in *Syngap1^+/c.3583-9G>A^* mice (t(2) = –5.71, *p* = 0. 0293, n = 3, left) but not controls (t(2) = –0.24, *p* = 0. 836, n = 3, right). (E) Representative Western blot of phosphorylated extracellular signal-regulated kinases 1 and 2 (p-ERK1/2) (∼42 and 44kDa) relative to total ERK1/2 expression (bottom) in HPC of *Syngap1^+/c.3583-9G>A^* (+) and control (−) mice. (F) P-ERK1/2 relative to total ERK1/2 expression in HPC of *Syngap1^+/c.3583-9G>A^*and control mice. Welch two sample t-test showed significant up-regulation of ERK1/2 signaling in *Syngap1^+/c.3583-9G>A^* mice (t(9.8) = 2.32, *p* = 0.043, n = 6 per group). (G) Representative Western blots of p-ERK1/2 protein (∼42 and 44 kDa) expression in HPC of *Syngap1^+/c.3583-9G>A^* (left) and control (right) mice, which received contralateral injections of either nt control dCas9-VPR (nt) or Syngap1 g44-dCas9-VPR (g44 in red). Shown relative to total ERK1/2 expression probed on the same membrane (bottom). (H) P-ERK1/2 relative to total ERK1/2 expression in HPC after injection of lentiviral vectors expressing either nt control dCas9-VPR or Syngap1 g44-dCas9-VPR (red). Paired t-test showed significant down-regulation of p-ERK1/2 in *Syngap1^+/c.3583-9G>A^* mice (t(3) = 3.24, *p* = 0.0478, n = 4, left) and controls (t(4) = 4.17, *p* = 0.014, n = 5, right). (I) Schematic representation of spontaneous alternations in Y-maze paradigm. (J) Number of arm entries in 8 minutes (min) for control or *Syngap1^+/c.3583-9G>A^*mice. Two-tailed t-test showed significantly more arm entries for *Syngap1^+/c.3583-9G>A^*mice (t(17) = –5.23, *p* = 6.827 × 10^-5^, n = 19). (K) Percentage of spontaneous alternations in 8 min for control or *Syngap1^+/c.3583-9G>A^*mice. Two-tailed t-test showed significantly reduced spontaneous alternations for *Syngap1^+/c.3583-9G>A^* mice (t(17) = 2.21, *p* = 0.0413, n = 19). (L) Schematic representation of experimental timeline. Mice received bi-lateral HPC injections of either nt control dCas9-VPR or Syngap1 g44-dCas9-VPR (red). 14 days post-surgery, mice were handled for three days before being subjected to Y-maze behavioral testing for 8 min. (M) Number of arm entries in 8 min for *Syngap1^+/c.3583-9G>A^* (n = 21) and control (n = 21) mice after injection of nt control dCas9-VPR or Syngap1 g44-dCas9-VPR (red). Syngap1 g44-dCas9-VPR did not rescue hyper-locomotor behavior. Two-way ANOVA showed significant effects for genotype (F(1) = 25.19, *p* = 1.25 x 10^-5^). There was no main effect of treatment (F(1, 38) = 0.75, *p* = 0.393), nor a treatment:genotype interaction effect (F(1, 38) = 1.76, *p* = 0.192). Data were pooled from three independent cohorts, with no cohort effect observed (F(2, 37) = 1.59, *p* = 0.218) (N) Percentage of spontaneous alternations in 8 min for *Syngap1^+/c.3583-9G>A^* (n = 21) and control (n = 21) mice after HPC injection of nt control dCas9-VPR or Syngap1 g44-dCas9-VPR (red). Syngap1 g44-dCas9-VPR rescued working memory deficits in *Syngap1^+/c.3583-9G>A^* mice but had no effect in control mice: Two-way ANOVA of genotype and treatment showed an interaction effect (F(1, 38) = 8.96, *p* = 0.0048). Post-hoc TukeyHSD revealed no significant difference between *Syngap1^+/c.3583-9G>A^*mice treated with Syngap1 g44-dCas9-VPR and control mice treated with nt control dCas9-VPR (difference = 1.90, 95% CI [–4.91, 8.71], p = 0.877), and a trend toward improved performance in *Syngap1^+/c.3583-9G>A^* mice treated with Syngap1 g44-dCas9-VPR compared to nt control group (difference = 6.93, 95% CI [–0.05, 13.92], p = 0.052). Data were pooled from three independent cohorts, with no cohort effect observed (F(2, 37) = 0.13, *p* = 0.881). Bars represent mean ± SEM error bars, * p < 0.05.

SynGAP is a Ras GTPase-activating protein and a negative regulator of Ras-extracellular signal-regulated kinase (ERK) signaling in neurons^31,40^. To investigate whether our CRISPRa strategy affects this downstream mechanism of SynGAP, we first confirmed that phosphorylated ERK1/2 (p-ERK1/2) is increased in HPC of *Syngap1^+/c.3583-9G>A^* mice compared to controls (**Fig. 3E & F**). Next, we performed stereotaxic surgery with contralateral HPC injections as above and found downregulation of p-ERK1/2 over total ERK1/2 due to Syngap1 g44-dCas9-VPR in both *Syngap1^+/c.3583-9G>A^* and control mice 14 days post-transfection (**Fig. 3G & H**). This data confirms that *Syngap1* CRISPRa can functionally suppress Ras-ERK signaling *in vivo*.

### Syngap1 CRISPRa rescues SRID-related behavior

Lastly, we aimed to examine whether *Syngap1* CRISPRa can rescue SRID-associated behaviors in *Syngap1^+/c.3583-9G>A^* mice. To address this, we first performed open field behavioral testing in adult mixed-sex mice. Consistent with previous reports^28,31^, *Syngap1^+/c.3583-9G>A^*mice traveled greater distances at higher speed compared to controls (**Extended Data Fig. 2A-C**), confirming a hyper-locomotor phenotype^27^. We did not detect an anxiety-like phenotype in this assay (**Extended Data Fig. 2D-E**). We next assessed spatial learning and memory using Barnes maze behavioral testing (**Extended Data Fig. 2F**). *Syngap1^+/c.3583-9G>A^*mice spent less time in the target quadrant and around the target hole (**Extended Data Fig. 2G-H**), associated with the cue during the acquisition phase, indicating spatial memory deficits. However, latency to first entry into the target hole did not differ between groups (**Extended Data Fig. 2I**). This likely reflects the hyper-locomotor phenotype in *Syngap1^+/c.3583-9G>A^* mice, which masks the readout for spatial memory performance in the Barnes maze (**Extended Data Fig. 2J**).

We next performed Y-maze behavioral testing in adult mixed-sex mice, which provides a readout of spatial working memory unconfounded by locomotor activity^44^. We replicated behaviors associated with this mouse model^31^: *Syngap1^+/c.3583-9G>A^* mice showed an increased number of arm entries compared to controls (**Fig. 3J**), confirming hyper-locomotor behavior. Further, we showed decreased alternative alternations in Syngap1*^+/c.3583-9G>A^* mice (**Fig. 3I & K**), indicating spatial working memory deficits^45^. To test the effects of Syngap1 CRISPRa on these behaviors, we performed bi-lateral HPC injections of either nt control dCas9-VPR or Syngap1 g44-dCas9-VPR and repeated Y-maze testing. Data were pooled from three independent cohorts with no significant cohort effects observed (**3M & N**). Syngap1 g44-dCas9-VPR did not rescue hyper-locomotor behavior (**Fig. 3M**). However, Syngap1 g44-dCas9-VPR rescued the spatial working memory deficit in *Syngap1^+/c.3583-9G>A^*mice, as indicated by a genotype:treatment interaction effect in spontaneous alternation performance (F(1, 38) = 8.96, *p* = 0.0048; **Fig. 3N**). Post-hoc Tukey’s HSD revealed that CRISPRa treatment did not have an effect on spontaneous alternations in control mice, consistent with data showing no CRISPRa-mediated upregulation of *Syngap1* mRNA and protein in this group. Further, *Syngap1^+/c.3583-9G>A^* mice treated with nt control showed significantly reduced spontaneous alternations compared to control mice treated with nt control (TukeyHSD: *p* = 0.0084), consistent with a spatial working memory deficit. In contrast, *Syngap1^+/c.3583-9G>A^*mice treated with Syngap1 g44-dCas9-VPR did not differ from nt control-treated control mice (TukeyHSD: *p* = 0.882), indicating CRISPRa rescue of the working memory deficit. In sum, our dual-lentiviral Syngap1 g44-dCas9-VPR approach successfully increased *Syngap1* mRNA and SynGAP protein expression, attenuated Ras-ERK1/2 signaling, and rescued spatial working memory deficits in *Syngap1^+/c.3583-9G>A^* mice. These findings highlight the therapeutic potential of CRISPR-mediated *SYNGAP1* gene activation for treating SynGAP haploinsufficiency.

### *SYNGAP1* CRISPRa activates human *SYNGAP1* in haploinsufficient patient neurons

Having developed *Syngap1* CRISPRa in a mouse model of SRID, we next tested whether our system can be translated to human and whether it is effective against other *SYNGAP1* loss-of-function mutations leading to SynGAP haploinsufficiency. We designed a gRNA library targeting regions near the human *SYNGAP1* TSS enriched for permissive hPTM H3K27ac (histone H3 lysine 27 acetylation) (**Fig. 4A)**. The gRNAs targeted the dCas9-VPR system to sites both up-stream (−1036, –424 bp) and down-stream (+29, +41 bp) of the TSS of human *SYNGAP1* (**Fig. 4B; Supplementary Table 1**). We packaged each construct into lentiviral vectors and screened the library in SH-SY5 cells. We used a nt gRNA with no homology to the human genome combined with dCas9-VPR as a control. Three days after treatment, SYNGAP1 g41-dCas9-VPR and SYNGAP1 g-1036-dCas9-VPR activated *SYNGAP1* mRNA (**Fig. 4C-D**). SYNGAP1 g-1036-dCas9-VPR upregulated *SYNGAP1* expression by approximately 2-fold, compared to a ∼1.5-fold increase with g41-dCas9-VPR. Notably, SYNGAP1 g41 is identical to the mouse homolog Syngap1 g44 (**Fig. 2C**), whereas SYNGAP1 g-1036 is unique to humans (**Supplementary Table 1**), highlighting the importance of using both preclinical mouse and human SRID models.

**Figure 4.**
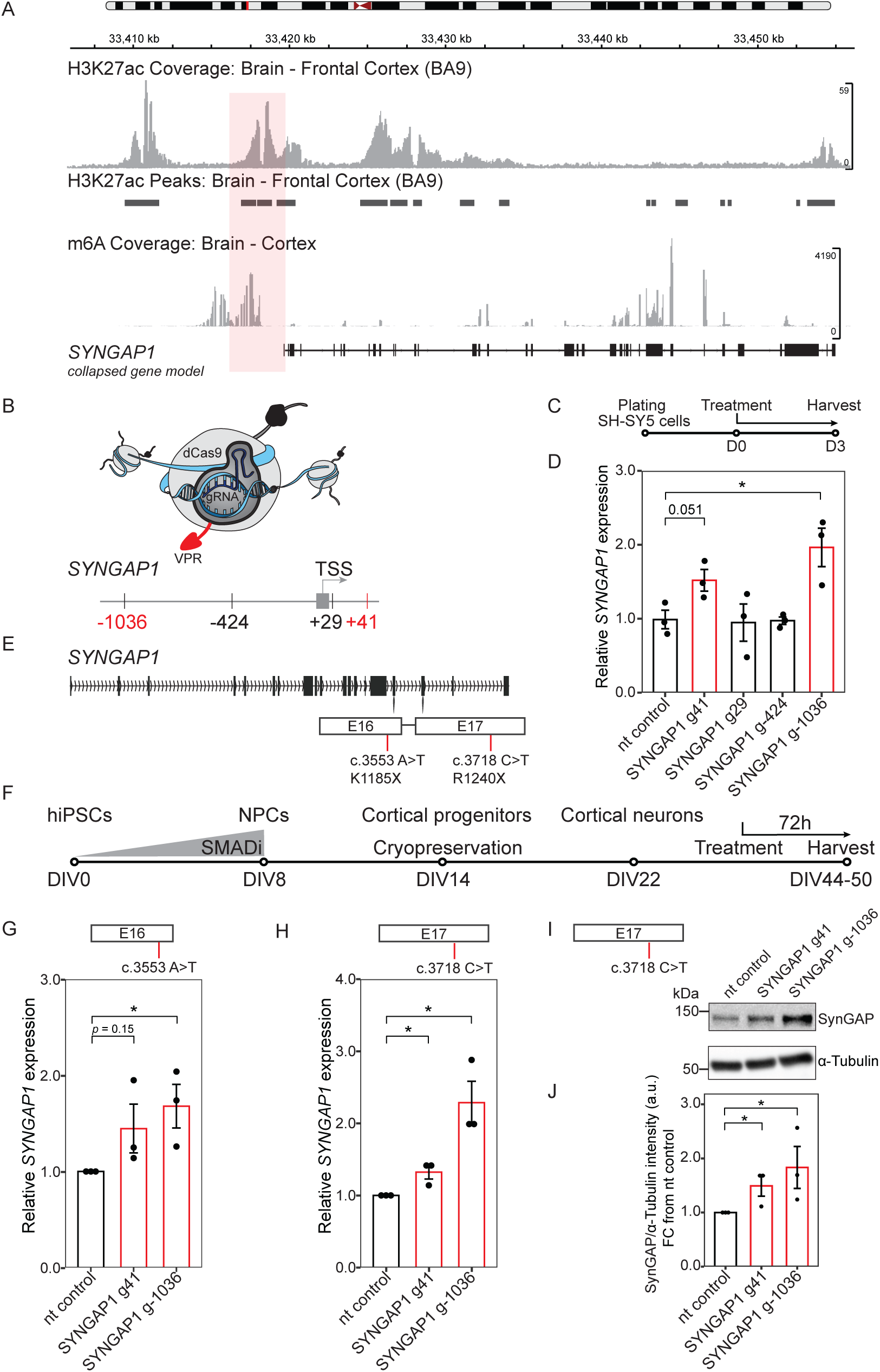
SYNGAP1 CRISPRa activates human *SYNGAP1* in haploinsufficient patient neurons. (A) Genome browser view of gene activation-associated histone mark H3K27ac enrichment near *SYNGAP1* transcription start site (TSS) in frontal cortex. Data from the Broad Institute’s Adult Genotype Tissue Expression (GTEx) Project. The red box highlights the locus targeted for SYNGAP1 CRISPRa gRNA design. (B) Illustration of SYNGAP1 CRISPRa tool and gRNA library up-(−1036 and –424 base pairs (bp)) and down-stream (29 and 41 bp) of the TSS of human *SYNGAP1*. (C) Timeline of lentiviral vector treatment of SH-SY5 cells. (D) *SYNGAP1* mRNA levels (RT-qPCR) in SH-SY5 cells (n = 3) treated with lentiviral vectors expressing CRISPRa non-targeting (nt) control and targeted at *SYNGAP1* locus (Syngap1 g-1036, g-424, g29 and g41). *SYNGAP1* mRNA expression is shown relative to nt control. Two-tailed t-test showed up-regulation of *SYNGAP1* mRNA for SYNGAP1 g-1036 (t(4) = 3.38, p = 0.0279) and trending up-regulation for SYNGAP1 g41 (t(4) = 2.76, p = 0.0510) compared to nt control. (E) Illustration of *SYNGAP1* loss-of-function mutations in exon 16 (c.3553A>T, K1185X) and exon 17 (c.3718C>T, R1240X) leading to SynGAP1 haploinsufficiency. Human induced pluripotent stem cell (hiPSC) lines carrying these mutations were used for analysis. (F) Time course of neuronal differentiation protocol of patient-derived hiPSCs and lentiviral vector CRISPRa treatment of hiPSC-derived neurons. (G) *SYNGAP1* mRNA expression (RT-qPCR) in day *in vitro* (DIV) 44 neurons differentiated from patient-derived hiPSC-line c.3553A>T (K1185X) after lentiviral vector transduction of either nt control dCas9-VPR, SYNGAP1 g41-dCas9-VPR or SYNGAP1 g-1036-dCas9-VPR (red). *SYNGAP1* mRNA expression is shown relative to nt control within same differentiation. Two-tailed t-test showed significant up-regulation of *SYNGAP1* in SYNGAP1 g-1036-dCas9-VPR (t(4) = 2.99, *p* = 0.040, n = 3), but not in SYNGAP1 g-41-dCas9-VPR conditions (t(4) = 1.76, *p* = 0.153, n = 3). (H) *SYNGAP1* mRNA expression (RT-qPCR) in day *in vitro* (DIV) 50 neurons differentiated from patient-derived hiPSC-line c.3718C>T (R1240X) after lentiviral vector transduction of either nt control dCas9-VPR, SYNGAP1 g41-dCas9-VPR or SYNGAP1 g-1036-dCas9-VPR (red). *SYNGAP1* mRNA expression is shown relative to nt control within same differentiation. Two-tailed t-test showed significant up-regulation of *SYNGAP1* in SYNGAP1 g-41-dCas9-VPR (t(4) = 3.39, *p* = 0.027, n = 3) and SYNGAP1 g-1036-dCas9-VPR (t(4) = 4.32, *p* = 0.012, n = 3). (I) Representative Western blot of SynGAP protein (∼150kDa) levels in haploinsufficient patient neurons (c.3718C>T, R1240X) treated with lentiviral vector nt control dCas9-VPR, SYNGAP1 g41-dCas9-VPR, or SYNGAP1 g-1036-dCas9-VPR. Loading control α-Tubulin (∼55kDa) probed on the same membrane (bottom). (J) SynGAP protein expression in haploinsufficient patient neurons (c.3718C>T, R1240X) treated with lentiviral vector nt control dCas9-VPR, SYNGAP1 g41-dCas9-VPR (red), or SYNGAP1 g-1036-dCas9-VPR (red). SynGAP expression is shown relative to nt control within same differentiation. One-tailed t-test showed significant up-regulation of SynGAP after SYNGAP1 g41-dCas9-VPR (t(4) = 2.59, *p* = 0. 030, n = 3) and SYNGAP1 g-1036-dCas9-VPR (t(4) = 2.14, *p* = 0. 049, n = 3) transduction. Bars represent mean ± SEM error bars, * p < 0.05.

To assess whether human *SYNGAP1* CRISPRa activated *SYNGAP1* in neurons, we tested SYNGAP1 g41-dCas9-VPR and g-1036-dCas9-VPR in primary cortical neurons dissected from hybrid humanized *Syngap1* mice. This previously validated mouse model was genetically modified to replace both mouse alleles for *Syngap1* with the human gene (*Syngap1*^Hu/Hu^) and was crossed with a *Syngap1* haploinsufficiency mouse model (*Syngap1*^+/-^) to generate *Syngap1* humanized haploinsufficient mice (*Syngap1*^Hu/-^, and *Syngap1*^Hu/+^ controls)^46^. After confirming *SYNGAP1* haploinsufficiency in *Syngap1*^Hu/-^ cortical neurons (**Extended Data Fig. 3A-C**), we transduced DIV14 neurons with either SYNGAP1 g41-dCas9-VPR, g-1036-dCas9-VPR or nt control dCas9-VPR. 7 days post-transduction, we harvested the cells and tested *SYNGAP1* mRNA expression (**Extended Data Fig. 3A**). Interestingly, *SYNGAP1* CRISPRa activated *SYNGAP1* only in haploinsufficient *Syngap1*^Hu/-^, but not in *Syngap1*^Hu/-^ neurons (**Extended Data Fig. 3D**), consistent with our observations in *Syngap1^+/c.3583-9G>A^* and control mice and with previous reports of CRISPRa targeting other dosage-sensitive genes^42,43^. Furthermore, contrary to our findings in SH-SY5 cells, SYNGAP1 g41-dCas9-VPR induced more robust *SYNGAP1* upregulation than SYNGAP1 g-1036-dCas9-VPR (**Extended Data Fig. 3D**), likely reflecting mouse-specific gene regulatory networks in *Syngap1*^Hu/-^ neurons.

Finally, we tested *SYNGAP1* CRISPRa in cortical excitatory neurons derived from human induced pluripotent stem cell (hiPSC) lines from two SRID patients carrying heterozygous loss-of-function mutations in exon 16 (c.3553A>T, K1185X) or exon 17 (c.3718C>T, R1240X) (**Fig. 4E**), both leading to haploinsufficiency. We previously characterized and confirmed *SYNGAP1* haploinsufficiency in these lines^47^. We allowed neurons to mature to DIV41 and treated with lentiviral vectors carrying SYNGAP1 g41-dCas9-VPR, SYNGAP1 g-1036-dCas9-VPR or nt control dCas9-VPR. At DIV44, SYNGAP1 g41-dCas9-VPR did not activate *SYNGAP1* mRNA in patient line-derived neurons, whereas SYNGAP1 g-1036-dCas9-VPR increased *SYNGAP1* mRNA in neurons derived from both lines (**Fig. 4G, Extended Data Fig. 3E**). To test CRISPRa activity at a later stage of neuronal development, we matured c.3718C>T neurons until DIV47 before treatment and collected mRNA three days later. At DIV50, both gRNAs activated *SYNGAP1*, with SYNGAP1 g41 showing modest (∼1.3-fold) and SYNGAP1 g-1036 stronger ∼2-fold induction (**Fig. 4H**). Lastly, protein analysis in c.3718C>T neurons revealed modest SynGAP upregulation with SYNGAP1 g41-dCas9-VPR and ∼2-fold upregulation with SYNGAP1 g-1036-dCas9-VPR (**Fig. 4I–J**). Taken together, these results confirm that we developed an efficient strategy to activate *SYNGAP1* in haploinsufficient neurons, demonstrate translatability to human models across different SRID mutations.

## Discussion

SRID is a severe NDD with no available treatment targeting its underlying genetic cause^7^. Here, we used an SRID mouse model with a pathogenic *SYNGAP1* variant and confirmed SynGAP haploinsufficiency with associated ERK pathway overactivation and working memory deficits^31^. Using this preclinical mouse model, we aimed to lay the groundwork for developing an effective therapy for SRID by establishing a CRISPRa strategy targeting the *Syngap1* locus. We showed that *Syngap1* CRISPRa activates *Syngap1* mRNA and protein, which restored ERK signaling downstream of SynGAP and rescued working memory deficits in the SRID model. Importantly, this strategy also effectively increased human *SYNGAP1* expression in SRID patient-derived excitatory cortical neurons with distinct *SYNGAP1* loss-of-function variants, demonstrating its adaptability and mutation-independent therapeutic potential.

In line with previous evidence that SRID results from *SYNGAP1* loss-of-function^5,31^, our findings in the *Syngap1^+/c.3583-9G>A^*mouse model confirm reduced *Syngap1* mRNA and protein. Furthermore, the expression of the pathogenic *Syngap1^+/c.3583-9G>A^* isoform correlated with reduced *Syngap1* mRNA expression. Hence, the cryptic splice acceptor introduced by this pathogenic variant seems to be more dominant over the canonical splice sites, leading to a premature stop codon in exon 17. This supports the notion of NMD as a pathogenic mechanism in SRID and underlines the potential of therapeutic intervention targeted at the functional allele. We identified reduced *Syngap1* expression across three brain regions, though the extent of protein reduction varied, with more pronounced decreases observed in the PFC and HPC compared to the NAc. Such region-specific differences may reflect intrinsic variation in *Syngap1* expression across neuronal populations and highlight potential regional vulnerabilities to SynGAP loss^3^. Interestingly, we observed marked within-population variability in *Syngap1* expression among control littermates that was notably reduced in *Syngap1^+/c.3583-9G>A^* mice. This aligns with the broader concept that gene expression variability contributes to developmental identity and functional plasticity, and its loss may constrain adaptive potential in disease states^48,49^. Importantly, this intrinsic variability has practical implications for developing therapeutic strategies aimed at *SYNGAP1* upregulation, as it can obscure treatment effects if not properly controlled. In our study, we mitigated this by employing within-animal controls and sufficiently powered behavioral cohorts, ensuring robust assessment of intervention efficacy.

SynGAP, a Ras-GTPase inhibitor, regulates synaptic strength by suppressing small GTPase signaling and limiting AMPAR incorporation through MAGUK PDZ domain binding^18,22^. Haploinsufficiency of SynGAP has major implications on synaptic function due to elevated basal GTPase activity, increased AMPAR insertion, and reduced LTP due to constitutively enhanced synaptic strength. Given the broad range of molecular downstream mechanisms affected by *Syngap1* loss-of-function, therapeutic approaches should target the primary deficit. Here, we developed a CRISPRa strategy directed at the *Syngap1* promoter region to selectively activate the functional allele in an SRID mouse model. Our approach achieves high specificity through promoter-targeted gRNA design, and we optimized its transcriptional efficacy by fusing a tri-partite activator to dCas9. Using CRISPRa, we successfully activated *Syngap1* in the HPC of SRID model mice and therefore, *Syngap1* mRNA and protein levels increased. Consequently, ERK hyperactivity was downregulated. As ERK signaling underlies hippocampal memory processes^50^, it is consistent that CRISPRa-mediated *Syngap1* upregulation also rescued working memory deficits in *Syngap1^+/c.3583-9G>A^* mice. However, Syngap1 CRISPRa did not rescue hyperactivity, which may be due to the targeted HPC administration in SRID mice. Previous work showed that genetic restoration of *Syngap1* in indirect pathway striatal projection neurons rescues motor phenotypes in *Syngap1^+/-^*mice^27^, supporting the importance of broader circuit-level interventions. Future therapeutic strategies could employ systemic or brain-wide delivery approaches, such as intracerebroventricular injection, intranasal, or intravenous administration with blood brain barrier-permeable vectors, to target all *Syngap1*-expressing brain regions.

When applying CRISPRa to restore expression of haploinsufficient genes, it is important to consider that excessive activation can negatively impact cellular health. This consideration is particularly relevant for genes with strict dosage requirements, for which promoter-directed upregulation may easily disrupt the delicate homeostasis of expression^51^. Interestingly, in our study, we observed *Syngap1* upregulation exclusively in HPCs of mutant mice but not in HPCs of control littermates, consistent with previous CRISPRa studies targeting other dosage-sensitive neurodevelopmental genes^42,43^. Several molecular mechanisms could account for this selective response. First, lower baseline expression in *Syngap1* haploinsufficient neurons may provide a greater dynamic range for CRISPRa, whereas transcription in wildtype neurons may already be near saturation^52^. Second, activation of the intact functional allele in haploinsufficient neurons may yield a pronounced effect without competing transcription from a second functional allele. Finally, tight homeostatic feedback in wildtype neurons may buffer against excessive *Syngap1* expression, preventing overactivation and potential toxicity^53^. Our and others’ findings indicate that CRISPRa of some genes occurs selectively in the haploinsufficient state^42,43^, underscoring its therapeutic promise while minimizing the risk of potential overexpression-related adverse effects.

An unexpected finding was the reduction of p-ERK levels in control mice following CRISPRa treatment. This may reflect a transient phase of SynGAP upregulation in wildtype neurons that was not captured at the timepoint analyzed (14 days post-surgery). Given that Ras-Raf-MEK-ERK signaling is a downstream target of SynGAP, normalization of this pathway may lag behind potential homeostatic mechanisms to prevent excessive activation of *SYNGAP1*. A temporal analysis will be required to determine whether CRISPRa initially upregulates *Syngap1* in control animals and subsequently stabilizes through such compensatory processes. Supporting this, our *in vitro* data show *Syngap1* activation seven days after transduction in control primary cortical neurons. CRISPRa-induced SynGAP upregulation may engage intrinsic regulatory mechanisms that fine-tune signaling homeostasis, emphasizing the importance of temporal precision when evaluating therapeutic gene activation *in vivo*.

Groundbreaking advances in CRISPR-based genome engineering have enabled the transition of this technology from the bench to the bedside, with CRISPR-Cas9 now approved by the FDA for treating sickle cell disease^34^. However, genome-editing approaches that rely on DNA cleavage are invasive and irreversible. In contrast, CRISPRa offers a safer and reversible alternative, making it an attractive therapeutic strategy. SRID is an ideal candidate for CRISPRa intervention, as it is a monogenic disorder and we and others have shown that restoring SynGAP protein levels can rescue SRID-associated phenotypes in mouse models^27,54^. Importantly, we demonstrate the translational potential of our approach by showing robust upregulation of *SYNGAP1* mRNA and protein in excitatory cortical neurons derived from SRID patient hiPSC lines. Notably, while a human analog of our highest performing mouse Syngap1 gRNA increased *SYNGAP1* expression, a human-specific gRNA achieved even greater activation, highlighting the adaptability of the system and importance of using both mouse and human models. These findings further support that CRISPRa acts in a mutation-independent manner and can thus be broadly applied across distinct loss-of-function variants. Integration with allele-specific silencing approaches, such as antisense oligonucleotides, could further enhance therapeutic efficacy^47,55^. Further, for therapeutic translation, future work will require development of a non-integrating adeno-associated virus (AAV) delivery platform optimized for brain-specific expression^56,57^. Overall, our results establish a promising framework for CRISPRa gene activation as a viable therapeutic approach for SRID.

## Materials and Methods

### Animals

*Syngap1^+/c.3583-9G>A^* mice and wildtype littermates were maintained on a mixed C57BL/6J and 129/SvEv background. Animals were housed under a 12 h light/dark cycle at a constant temperature (23 °C) with ad libitum access to food and water. All procedures were conducted in accordance with National Institute of Health guidelines and approved by the University of Pennsylvania Institutional Animal Care and Use Committee. The facility is accredited by the Association for Assessment and Accreditation of Laboratory Animal Care.

### CRISPR-dCas9 and gRNA construct design

gRNA constructs were designed with benchling targeting the upstream promoter and first intron of the mouse and human *Syngap1*/*SYNGAP1* locus (**Supplementary Table 1**). Efficiency score from^58^ was prioritized. Two separate plasmids were used, one for gRNA expression, driven by the ubiquitous U6 promoter (https://www.addgene.org/114199/), and one for dCas9-VPR expression, driven by the neuronal hSyn promoter (https://www.addgene.org/114196/).

Plasmid #114199 was digested with BbsI high fidelity restriction enzyme right after the U6 promoter. Two, 25-nucleotide complementary oligos with overhangs specific to the BbsI cut plasmid were ordered from IDT. Those oligos were annealed and phosphorylated with PNK, then ligated into the plasmid using T4 ligase. Those plasmids were then transformed into the lentiviral vector with STBI3 bacteria. Correct plasmid constructs were selected for as single colonies using ampicillin resistance and sequenced at the insertion site using Sanger sequencing to confirm.

Then each gRNA plasmid and the dCas9-VPR plasmids were prepped en masse for lentiviral packaging. 5 mL of culture tubes were grown with LB and ampicillin for 5 h and added to 1 L of LB for overnight growth. Plasmids were purified using the Qiagen Mega kit (#12381) following their quickstart protocol.

### Lentiviral vector production

Third-generation lentiviral vectors were produced using HEK293T cells. Cells were maintained in DMEM supplemented with 10% fetal bovine serum (FBS) and 1% penicillin–streptomycin (P/S) and seeded at ∼4.5 x 10^6 cells per T225 flask. Transfections were performed at ∼80% confluency using TransIT-Lenti reagent and plasmids encoding different CRISPRa constructs, pRRE, pRSV-Rev, and pVSV-G at established molar ratios. Cells were exposed to transfection medium containing 25 µM chloroquine for 5-6 h before being returned to harvesting medium (DMEM + 10% FBS + 1% P/S + 10 mM HEPES, pH 7.6). Supernatants were collected at 24, 48, and 72 h post-transfection, clarified through 0.45 μm filters, and concentrated by ultracentrifugation at 12,000 rpm for 20 h at 4°C. Viral pellets were resuspended in chilled formulation buffer and aliquoted for storage at –80 °C.

### Intra-HPC transduction

Mice were anesthetized with 5% isoflurane in oxygen (1 L/min) for induction and maintained at 1.5% isoflurane (0.5 L/min) during surgery. Ophthalmic ointment was applied to prevent corneal drying, and pedal reflexes were checked before incision. The surgical site was disinfected with 7.5% povidone-iodine and 70% ethanol, and meloxicam was administered for analgesia. A midline scalp incision (∼2 cm) was made to expose bregma and lambda, and skull coordinates were determined relative to bregma. The dorsal hippocampus (HPC) was targeted bilaterally ^59,60^ using the following stereotaxic coordinates: –2.2 mm (anterior/posterior), ±2 mm (medial/lateral), and –1.8 mm (dorsal/ventral) at a 7° angle from the midline. Small craniotomies were made with a dental drill. Lentiviral constructs were prepared at a 1:4 dilution with sterile saline solution. A total of 1.5 µL was injected per hemisphere at 0.2 μL/min followed by a 5 min rest period before needle withdrawal. Incisions were closed with 4.0 sutures, and mice were placed under a heat lamp until ambulatory. Post-surgical monitoring was performed daily for a week to assess recovery.

### Mouse behavioral testing

**Open field.** 26 3-4 months old mixed-sex mice (13 *Syngap1^+/c.3583-9G>A^*; 13 controls) were tested in the open field. The experimenter was blinded to genotype throughout the whole procedure. The open field test was conducted in an Open Field Arena (Data Sciences International, #760190) equipped with a camera (White Matter LLC e3v vision). Mice were transferred to the testing room and habituated for 1 h prior to testing. Each mouse was placed in the center of the arena and allowed to freely explore for 10 min. Behavior was video recorded using the *e3v* software. Following each trial, the arena was cleaned with 70% ethanol. Video recordings were scored using the ANYmaze software.

**Barnes Maze**. Eleven 11-week-old mixed-sex mice (5 *Syngap1^+/c.3583-9G>A^*, 6 control littermates) were tested in the Barnes Maze. The experimenter was blinded to genotype throughout the whole procedure. The Barnes Maze (San Diego Instruments; #7001-0235) consisted of a 36-inch diameter white circular platform with 20 holes (2-inch diameter), elevated 36 inches above the floor. All holes except the target were occluded using dummy box undersides. The target location remained constant for each mouse but was alternated between animals to control for environmental variables. Visual cues were positioned around the arena, with one directly behind the target location. Maze illumination was set to ∼1000 lux. Mice were handled for 3 days before testing with consistent PPE across handling, training, and testing. On testing days, animals were habituated to the procedure room for ∼30 min. Mice were placed in a start box at the center for 1 min before release. Each trial lasted up to 150 s; entry into the target box was followed by a 60 s stay. For failed trials, latency was recorded as 150 s and mice were gently guided into the target box. Between trials, the target box and maze surface were cleaned with 70% ethanol. Animals underwent two 150 s trials per day with 15-30 min inter-trial intervals. After 4 days of acquisition, a single 2-min probe trial with all holes occluded was performed on testing day. Video recordings of the testing day were scored using the ANYmaze software.

**Y-Maze.** To assess working memory, 19 mixed-sex mice aged 12-16.5 weeks (11 *Syngap1^+/c.3583-9G>A^*, 8 control littermates) were tested in the Y-maze spontaneous alternation task. To assess working memory after intra-HPC transduction of Syngap1 CRISPRa, three independent cohorts comprising a total of 42 surgerized mice (21 *Syngap1^+/c.3583-9G>A^*, 21 control littermates) were tested. Cohorts were pooled for analysis. The examiner was blinded to genotype and treatment throughout the procedure. The maze (Harvard Bioscience #760079) consisted of three identical arms (designated A, B, and C) arranged in a “Y” configuration. Mice were handled for 3 days before testing with consistent PPE across handling, and testing. On testing day, mice were habituated to the testing room for 30 min. Each trial began with the mouse placed at the distal end of arm C, and its behavior was recorded for 8 min. The maze was cleaned with 70% ethanol between trials to minimize odor cues. Arm entries were defined as the placement of all four paws into an arm. Total arm entries and the sequence of entries were recorded. Spontaneous alternation was defined as successive entries into three different arms (e.g., ABC, BCA, or CAB). The percentage of spontaneous alternation was calculated as:

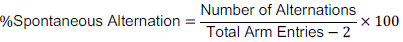

### N2A culture and transduction

Murine Neuro-2a (N2a) cells (ATCC CCL-131) were maintained in EMEM (ATCC #30-2003) supplemented with 10% fetal bovine serum (FBS; ATCC #30-2020). Cells were cultured at 37°C and 5% CO_2_ and passaged at 70-90% confluency using Trypsin/EDTA (ATCC #30-2101). Cryopreservation was performed in EMEM containing 10% FBS and 5% DMSO, with vials stored in liquid nitrogen. For treatment, cells were plated at ∼300,000 cells per well in a 6-well plate and transfected with lentiviral vectors the following day. Cells were 70-80% confluent at the time of infection; 1.5 μL of mixed dCas9-VPR (1 µL) and gRNA (0.5 µL) lentiviral vectors was added per 2 mL of fresh media (1:2 dilution). After 72 h, cells were examined for mCherry expression and lysed for downstream assays.

### SH-SY5Y culture and transduction

Human SH-SY5Y neuroblastoma cells (ATCC CRL-2266) were cultured in a 1:1 mixture of EMEM and DMEM/F12 supplemented with 10% FBS (ATCC #30-2020) at 37°C and 5% CO_2_. Cells were passaged at 70-90% confluency and cryopreserved in complete medium containing 5% DMSO. For lentiviral vector transduction, cells were seeded and treated as described above.

### Primary cortical neuron culture

Primary cortical neurons were prepared from embryonic day 16.5 (E16.5) *Syngap1^+/c.3583-9G>A^* and littermate control mouse embryos. Briefly, plates were pre-coated with poly-D-lysine (1 mg/mL) overnight, washed, and dried. Embryos were collected by CO_2_ euthanasia of pregnant dams followed by cervical dislocation and rapid embryo dissection in ice-cold PBS. Cortices were isolated under a dissecting microscope, meninges removed, and hippocampus/midbrain discarded to enrich for cortical tissue. Tissue was washed in OptiMEM + GlutaMAX (Gibco #51985-034), mechanically dissociated by trituration, and plated at a density of 2 brains per 6-well plate in OptiMEM + GlutaMAX (Gibco). After 2 h, medium was replaced with neuronal culture medium (Neurobasal supplemented with 2% B27 (Gibco), 0.5 mM GlutaMAX (Gibco #51985-034), and 1% Penicillin– Streptomycin (Gibco #15140-122). Cytosine β-D-arabinofuranoside (AraC, 0.5 µM) was added at 3 DIV to limit glial proliferation. Neurons were maintained at 37 °C, 5% CO_2_, with half-medium changes every other day. Cells were transduced with lentiviral vectors in concentrations as above on DIV4 and lysed on DIV11.

### Patient-derived hiPSC culture and excitatory cortical neuron differentiation

We previously described in detail the generation, validation and maintenance of the SRID patient-derived hiPSCs used in this study^47,61^. Briefly, hiPSCs were maintained under standard iPSC conditions, transitioned to feeder-free culture on human embryonic stem cell (hESC)-qualified Matrigel (Corning) in mTeSR1 medium (StemCell Technologies), and cryopreserved in 90% FBS/10% DMSO after ≥2 passages. Feeder-free stocks were expanded in mTeSR1 before differentiation.

Prior to differentiation, plates were coated with growth factor-reduced Matrigel (1 mg in 24 mL DMEM/F12; 2 mL per 35 mm well, incubated at 37°C for ≥1 h). HiPSCs were seeded in mTeSR1 at ∼50,000 cells/cm^2^. Once cultures reached ≥60% confluency, differentiation into neural progenitor cells (NPCs) was initiated with daily media changes containing SB431542 (10 µM; Tocris), LDN193189 (1 µM; Stemgent), IWR1 (1.5 µM; Tocris), and B27 without vitamin A (Invitrogen) ^62^. Cells were passaged with accutase (Invitrogen) on days 4 and 8 and replated onto coated plates with Y-27632 (10 µM) at ∼210,000 cells/cm^2^ (day 4) and ∼155,000 cells/cm^2^ (day 8). From days 8-14, NPCs were expanded in Neural Expansion Media (50% Advanced DMEM/F12, 50% Neurobasal, Neural Induction Supplement; Invitrogen) and subsequently cryopreserved in 90% FBS/10% DMSO.

NPCs were thawed from cryopreserved stocks and directed into differentiation. Cells were plated on GFR Matrigel-coated plates at ∼285,000/cm^2^ in N2B27(−) medium [2:1 DMEM/F12 (Invitrogen):Neurobasal (Invitrogen), 1/3x N2 (Invitrogen), 2/3x B27 without vitamin A (Invitrogen), 1x glutamine (Invitrogen), 50 μM β-mercaptoethanol (Invitrogen), and 25 ng/mL Activin A (BioTechne)], with daily media changes. After 6-9 days, when most cells showed neuronal morphology, cultures were passaged with 0.5 mM EDTA and replated at ∼155,000/cm^2^ on poly-D-lysine (10 μg/mL; Sigma) and GFR Matrigel-coated plates in N2B27(+) medium [same base medium with B27 + vitamin A, plus 20 ng/mL brain-derived neurotrophic factor (BDNF) and 20 ng/mL glial-derived neurotrophic factor (GDNF)]. Cultures were maintained with half-media changes three times per week until harvest. Neurons received lentiviral transduction at DIV 41 or DIV47 as described above and harvested for downstream analysis at DIV44/DIV50.

### Mouse brain tissue and cell lysate collection

For tissue collection, mice were sacrificed by cervical dislocation, and brains were placed in ice-cold phosphate-buffered saline containing protease inhibitors (Roche cOmplete #4693159001). One-millimeter coronal sections of the PFC, NAc, and HPC were prepared using a brain block, and regions were micro-dissected with a 2-mm punch (Harris) under a fluorescence stereoscope (Leica). For *in vivo* transduction studies, tissue was visually inspected to confirm mCherry expression of the plasmid for accurate HPC targeting and construct expression before punching. 2-mm punches were cryopreserved at –80 °C. Cells were washed briefly with PBS and lysates were collected in 500 µL of TRIzol or RLT for RNA isolation or in 150 µL of ice-cold RIPA buffer for protein isolation. Samples were either processed immediately or stored at –80 °C.

### Real-time qPCR

Total RNA was isolated using the RNeasy Mini Kit (Qiagen #74004) according to the manufacturer’s instructions, and 500 ng was reverse transcribed with Superscript IV (Bio-Rad #1708891). qPCR was performed on a QuantStudio 7 Flex Real-Time PCR System using TaqMan assays in 386-well plates. For mouse *Syngap1*, the following primers and probe were used: Forward 5’-GCGAGAAGAGTACAAGCTCAA; reverse 5’-CCTCTCCTTCAGCGAATGTATC; probe 5’-/56-FAM/TCCTCATAC/ZEN/TCCTTCACCCTGTCCAG/3IABkFQ. For human *SYNGAP1*, the following primers and probe were used: 5’-GAACGAAGTCACAACCCAAAC; reverse 5’-AGGACTCAGCAAGGACTCA; probe 5’-/56-FAM/TTCGGAAGC/ZEN/GAGGCAGGATCTG/3IABkFQ. Expression levels were normalized to *Actb*/*ACTB* (Mm.PT.58.29001744.g, Hs.PT.39a.22214847 IDT) in a multiplexed reaction, and relative mRNA abundance was calculated using the 2^-ΔΔCt^ method.

To detect the mutant *Syngap1^+/c.3583-9G>A^* transcript Power SYBR™ Green (Thermo Fisher Scientific #4367659) was used with the following primers: Forward 5’-CGGAGCGGACCGTAGCC, reverse 5’-CACCTGCCCGCCTGTCC.

### Western blotting

Frozen tissue punches were lysed in 150 µL RIPA buffer, homogenized with an electric pestle, incubated on ice for 10 min, and sonicated at high frequency for 10 s. Lysates were incubated on ice for an additional 5 min, centrifuged (14,000 g, 10 min, 4 °C), and the supernatant collected. Samples were aliquoted, quantified using a BCA assay (Thermo fisher scientific #23225), and stored at –80 °C or processed immediately. Proteins were separated on precast polyacrylamide gels (4-20% Mini-PROTEAN® TGX™, Bio-Rad #4561095) and transferred to PVDF membranes (0.2 μm; Bio-Rad #1620175) using wet transfer (250-350 mA, 120 min at 4 °C). Membranes were blocked in 5% bovine serum albumin (BSA; fisher scientific)/TBS-T for 60 min, incubated with primary antibodies [SynGAP (D20C7) Rabbit #5539 (Cell Signaling), α-Tubulin Rabbit #2114 (Cell Signaling), ATP5F1 #ab117991 (Abcam)] at 1:1000 overnight at 4 °C, and washed 3 x in TBS-T for 10 min each the following day. Secondary antibodies (1:10,000 in TBS-T) were applied for 60 min at room temperature, followed by 3 10-min washes in TBS-T. Detection was performed with ECL reagent and imaged using a chemiluminescence imager (Amersham 600RGB). Membranes were stripped (Restore™ Western Blot Stripping Buffer; Thermo Fisher Scientific #21059) and re-probed as needed.

### Statistical analysis

All statistical analyses were performed in RStudio (v2023.12.1+402). Data normality was assessed using the Shapiro-Wilk test. Normally distributed data were analyzed with parametric tests (two-way ANOVA with Tukey HSD post hoc corrections or Student’s *t*-tests, paired or unpaired as specified in figure legends). Non-normally distributed data were analyzed using the Wilcoxon rank-sum test. Outliers were identified, where appropriate, with the Grubbs test. Statistical significance was defined as *p* < 0.05. Data are presented as bar graphs with overlaid dot plots generated in ggplot2, with error bars representing the standard error of the mean (SEM).

## Acknowledgments

We thank the Neurobehavior Testing Core at the University of Pennsylvania, especially W. T. O’Brien and B. Ciesielski, for providing resources and training for animal behavioral testing. Further, we thank all members of the Heller lab at the University of Pennsylvania and all members of the Center for Epilepsy and Neurodevelopmental Disorders (ENDD) for ongoing discussion and feedback for this work. This study was funded by the National Institute of Health (R37 NS036715 to RLH), the ENDD, the SynGAP Research Fund (2020 to 2021 to R.L.H. and S.Z.), the STXBP1 Foundation (71923 to E.A.H.), the Orphan Disease Center (MDBR-23-008-SynGAP to E.A.H.), the European Union’s Horizon 2024 research and innovation programme under the Marie Skłodowska-Curie grant agreement (101210026 to L.S.), the Hartwell foundation (Hartwell Fellowship to M.B.R).

## Author Contributions

L.S., M.B.R., and E.A.H. designed research; L.S., M.B.R., S.A., S.Z., N.M., K.L.R.-A., M.H., K.S.C., J.W., E.A.W., L.V.D., I.H., Y.A., and R.J. performed research; L.S., M.B.R., K.L.R.-A., and N.M. analyzed data; R.L.H., G.P., D.L.F., B.L.D., B.L.P., and E.A.H. supervised the project. L.S. prepared the manuscript with input from M.B.R., and E.A.W. All authors discussed results and reviewed the manuscript.

## Extended Data

**Extended Data Fig. 1.**
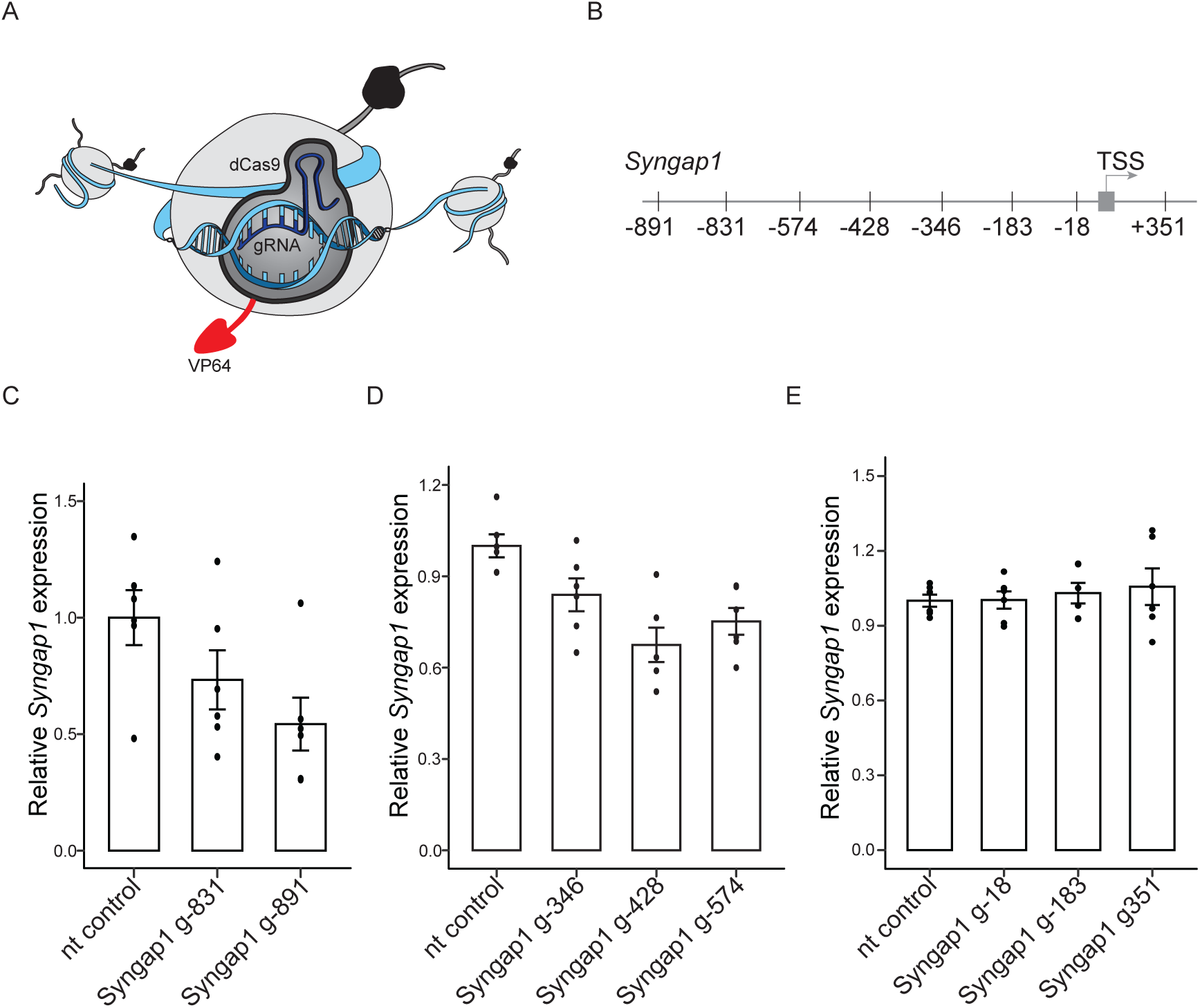
dCas9-VP64 targeted at *Syngap1* locus does not activate *Syngap1* in N2A cells. (A) Illustration of CRISPRa tool with dCas9 fused to the transcriptional activator VP64. (B) Illustration of gRNA library up-(−891, –831, –574, –428, –346, –183, and –18 base pairs (bp)) and down-stream (351 bp) of the TSS of mouse *Syngap1* screened with dCas9-VP64. (C-E) *Syngap1* mRNA levels (RT-qPCR) in N2a cells (n = 6) transduced with dCas9-VP64 and non-targeting (nt) control or targeting gRNAs (Syngap1 g-891 and g-831; g-346, g-428, and g-574; g-18, g-183, and g351). *Syngap1* mRNA expression is shown relative to nt control. *Syngap1* was not activated. Bars represent mean ± SEM error bars.

**Extended Data Fig. 2.**
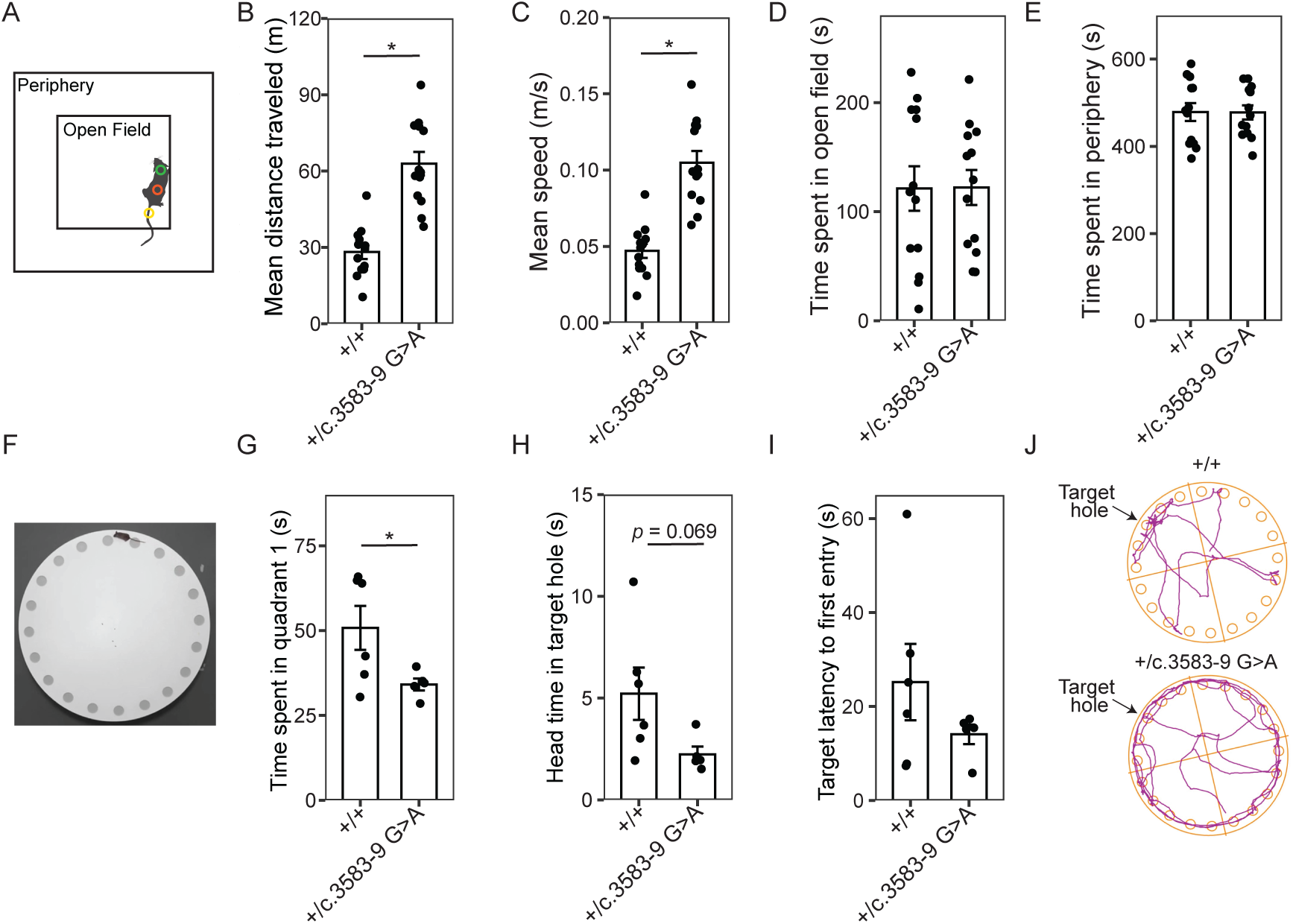
*Syngap1^+/c.3583-9G>A^* mice show hyper-locomotor behavior. (A) Illustration of open field paradigm. (B) Mean distance travelled in meters (m) in 10 minutes (min) for control or *Syngap1^+/c.3583-9G>A^* mice. Two-tailed t-test showed significantly greater distances for *Syngap1^+/c.3583-9G>A^* mice (t(24) = –6.46, *p* = 1.113 × 10^-6^, n = 26). (C) Mean speed in 10 minutes (min) for control or *Syngap1^+/c.3583-9G>A^* mice. Two-tailed t-test showed *Syngap1^+/c.3583-9G>A^* mice were significantly faster than controls (t(24) = –6.47, *p* = 1.089 × 10^-6^, n = 26). (D) No significant differences were observed between control or *Syngap1^+/c.3583-9G>A^*mice in time spent in the open field. (E) No significant differences were observed between control or *Syngap1^+/c.3583-9G>A^*mice in time spent in the peripheral zone. (F) Image of Barnes maze arena. (G) Time spent (s) in quadrant 1 (Q1) in 2 min on testing day. Mice learnt to associate the target hole with a cue during acquisition phase, which was situated in the center of Q1. *Syngap1^+/c.3583-9G>A^* mice spent significantly less time in Q1 (t(9) = 2.27, *p* = 0.049, n = 11), indicating spatial memory deficits. (H) Time of mouse head entering target hole during 2-min testing period. *Syngap1^+/c.3583-9G>A^* mice spent less time in target hole (t(5.8) = 2.22, *p* = 0.069, n = 11), indicating spatial memory deficits. (I) Target latency to first entry on testing day. No significant group differences were detected (t(9) = 1.21, *p* = 0.257, n = 11). (J) Barnes maze tracking plots of control (top) and *Syngap1^+/c.3583-9G>A^* (bottom) mouse showing hyper-locomotor phenotype. Bars represent mean ± SEM error bars, * p < 0.05.

**Extended Data Fig. 3.**
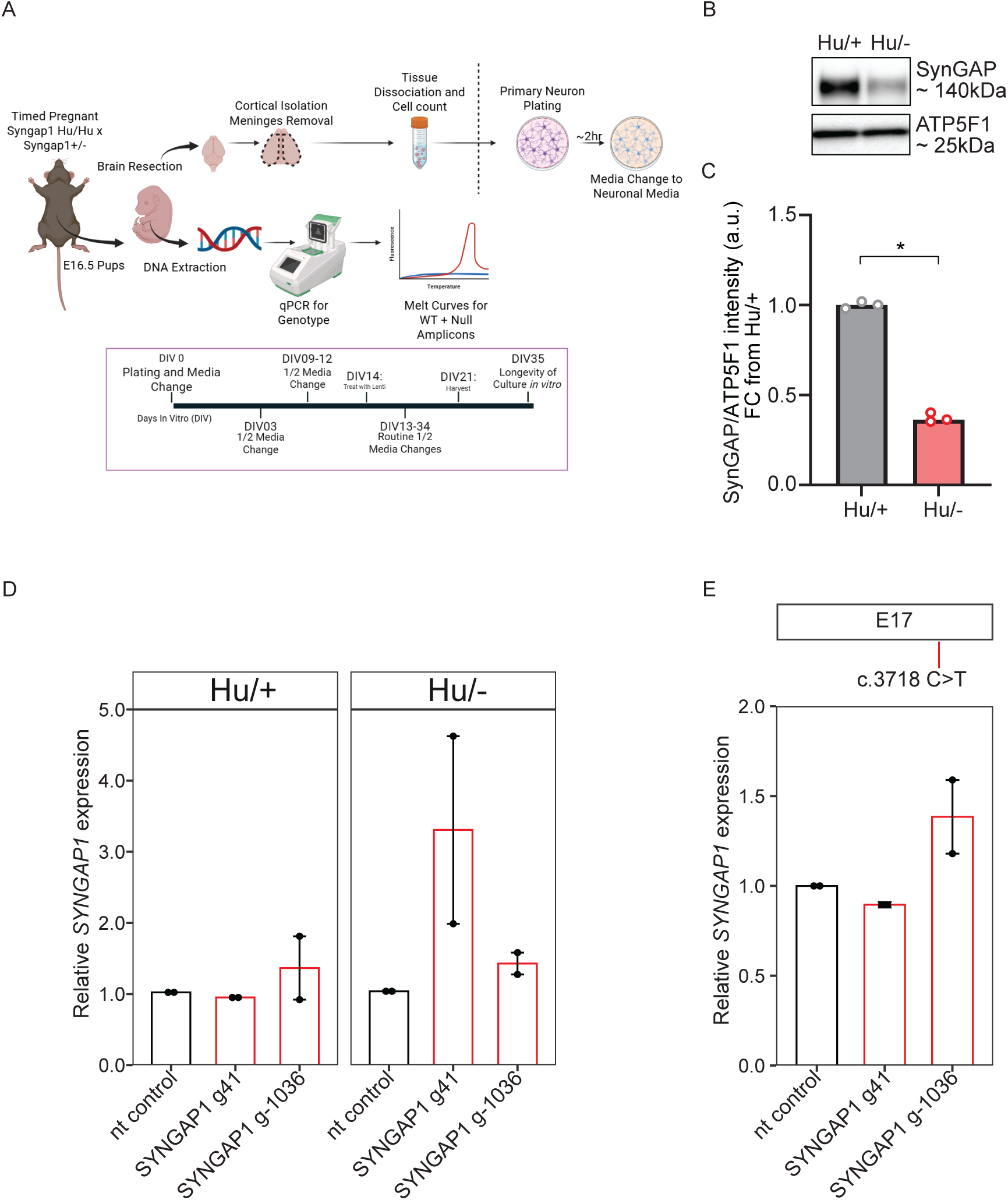
SYNGAP1 CRISPRa activates human *SYNGAP1* in haploinsufficient humanized mouse cortical neurons. (A) Experimental timeline of cortical neuron dissection from hybrid humanized *Syngap1*^Hu/+^ and *SYNGAP1* haploinsufficient Syngap*1*^Hu/-^ mice. (B) Representative Western blot showing SynGAP (∼150kDa) haploinsufficiency in Syngap*1*^Hu/-^ cortical neurons at day in vitro (DIV) 12. Loading control ATP15F1 (∼25kDa) probed on the same membrane (bottom). (C) SynGAP protein expression in haploinsufficient Syngap*1*^Hu/-^ cortical neurons. SynGAP expression is shown relative to *Syngap1*^Hu/+^. Two-tailed t-test showed significant reduction of SynGAP in Syngap*1*^Hu/-^ DIV12 cortical neurons (t(4) = 33.64, *p* <0.0001, n = 3). (D) *SYNGAP1* mRNA expression (RT-qPCR) in DIV21 neurons dissected from Syngap*1*^Hu/+^ (left) or Syngap*1*^Hu/-^ (right) mouse cortices after 7-day lentiviral vector transduction of either nt control dCas9-VPR, SYNGAP1 g41-dCas9-VPR or SYNGAP1 g-1036-dCas9-VPR (red). *SYNGAP1* mRNA expression is shown relative to nt control. Data points represent average of 1-4 technical replicates from two dissections (n = 2 biological replicates. Cortices of fetuses from each genotype were pooled prior to plating. (E) *SYNGAP1* mRNA expression (RT-qPCR) in DIV44 neurons differentiated from patient-derived hiPSC-line c.3718C>T (R1240X) after lentiviral vector transduction of either nt control dCas9-VPR, SYNGAP1 g41-dCas9-VPR (red) or SYNGAP1 g-1036-dCas9-VPR (red). *SYNGAP1* mRNA expression is shown relative to nt control within same differentiation. N = 2 biological replicates. Bars represent mean ± SEM error bars, * p < 0.05.

