## Supplemental Table 1 for "CRISPR-mediated transcriptional activation as a mutation-independent therapeutic strategy for *SYNGAP1*-related intellectual disability"

Tables

**Supplementary Table 1. Guide RNA (gRNA) sequences used in this study.**

| dCas9-VPR | | | | | | |
|---|---|---|---|---|---|---|
| ID | Strand | Sequence | PAM | Specificity Score | Efficiency Score | Species |
| SYNGAP1 g-1036 | 1 | GTCAGGGTTATACCTCTCGG | AGG | 91.2743603 | 71.89596303 | Human |
| SYNGAP1 g-424 | 1 | TGTCTCAAATTACCCCACAG | TGG | 68.4799745 | 77.60193043 | Human |
| SYNGAP1 g29 | 1 | TCTCGAGCCTCCATCCATCG | GGG | 83.0109294 | 64.34380838 | Human |
| SYNGAP1 g41 | -1 | AGGGGGCATAGGACATCGCG | GGG | 91.3941225 | 72.70699142 | Human |
| Syngap1 g-572 | -1 | TGTTAATCTTATTCGAACCC | TGG | 88.3670411 | 58.18400875 | Mouse |
| Syngap1 g-288 | -1 | TACCCAGGGAGACCCGGAAA | AGG | 76.3472041 | 50.90541354 | Mouse |
| Syngap1 g2 | -1 | AGGCTCGAGACCTGCTCATC | AGG | 78.577409 | 43.18387708 | Mouse |
| Syngap1 g44 | -1 | AGGGGGCATAGGACATCGCG | GGG | 92.4727299 | 72.70699142 | Mouse |

| dCas9-VP64 | | | | | | |
|---|---|---|---|---|---|---|
| ID | Strand | Sequence | PAM | Specificity Score | Efficiency Score | Species |
| Syngap1 g-891 | -1 | CCTAGGAAGGGCCCAAATAT | TGG | 78.68 | 28.78 | Mouse |
| Syngap1 g-831 | -1 | TGAGTTAGTGACCGATTGCG | AGG | 96.65 | 62.89 | Mouse |
| Syngap1 g-574 | -1 | TGTTAATCTTATTCGAACCC | TGG | 88.37 | 58.18 | Mouse |
| Syngap1 g-428 | -1 | CACCTCCATACTCAACAAAT | TGG | 67.33 | 31.76 | Mouse |
| Syngap1 g-346 | 1 | CAGCTTCTACGGGACTTCCC | TGG | 78.31 | 51.34 | Mouse |
| Syngap1 g-183 | 1 | TCGCGCTGTCTCCGGGCGAC | GGG | 95.90 | 34.50 | Mouse |
| Syngap1 g-18 | -1 | CTCGCGCCGCCCGACGCGGT | GGG | 98.20 | 29.30 | Mouse |
| Syngap1 g351 | 1 | TTCCTGCTTGTGACCGCGCG | TGG | 98.00 | 46.10 | Mouse |
